# UniverSC: a flexible cross-platform single-cell data processing pipeline

**DOI:** 10.1101/2021.01.19.427209

**Authors:** S. Thomas Kelly, Kai Battenberg, Nicola A. Hetherington, Makoto Hayashi, Aki Minoda

## Abstract

Single-cell RNA-sequencing analysis to quantify RNA molecules in individual cells has become popular owing to the large amount of information one can obtain from each experiment. We have developed UniverSC (https://github.com/minoda-lab/universc), a universal single-cell processing tool that supports any UMI-based platform. Our command-line tool enables consistent and comprehensive integration, comparison, and evaluation across data generated from a wide range of platforms.

## Introduction

Single-cell genomics technologies have led to a recent surge in studies of cellular heterogeneity. An important step in single-cell RNA-sequencing (scRNA-seq) is adding a unique molecular identifier (UMI) to each RNA molecule early on during library preparation to reduce the effect of amplification bias after capturing individual cells either in gel emulsion with beads or in wells. Recently developed technologies can perform scRNA-seq for thousands of cells and some are commercially available (e.g. Chromium by 10x Genomics, Nadia by Dolomite Bio, and ddSEQ by Bio-Rad). scRNA-seq is being applied to a whole range of tissues as well as whole organisms (Cao et al., 2017; Regev et al., 2017; The Tabula Muris Consortium, 2018). It is expected for scRNA-seq to become more accurate, more reliable, and with lower cost per cell, being feasible for a wide range of studies as the technology is developed further (Kulkarni *et al*., 2019).

Chromium platform (10x Genomics; Zheng et al., 2017) has gained a significant market-share in single-cell genomics, in part due to their user-friendly end-to-end workflows spanning from experimental kits to bioinformatics data processing tools. However, there are many other technologies including DropSeq (Macosko et al., 2015), Nadia (Dolomite Bio; Macosko et al., 2015), ICELL8 (Takara Bio; Goldstein et al., 2017), inDrop (Veres & Lee, 2018), SmartSeq3 (Hagemann-Jensen et al., 2019), and Quartz-seq (Sasagawa et al., 2013). Each single-cell platform has different advantages, and some may be more suitable than others for different sample types, especially those with challenging cell properties such as large cells. Therefore, an optimal platform should be selected from those available for the sample type of interest. This may be more critical for tissue samples and non-model organisms since many commercially available systems are designed for mammalian cells. When evaluating different platforms, different data processing pipelines are often used that have been specifically developed for each platform (Cell Ranger: 10x Genomics, 2020; InDrops: Veres & Lee, 2018; ddSeeker: Romagnoli et al., 2018). However, they may have different data processing criteria, making comparisons across platforms unreliable and challenging.

Leveraging on the fact that most scRNA-seq technologies use similar systems of cell barcodes and UMIs, we developed UniverSC, a shell utility that works as a wrapper for Cell Ranger, which can handle datasets generated by a wide range of single-cell technologies. Cell Ranger was chosen as a unifying pipeline for several reasons: 1) it is optimized to run in parallel on a cluster and the installation is fairly straightforward, 2) many labs working on single-cell analysis are likely to already be familiar with the outputs, 3) many tools have already been released for downstream analysis of the output format due to its popularity, 4) the rich summary information and post-processing is useful for further optimization and troubleshooting if necessary, and 5) the latest open-source release (version 3.0.2) has been optimized further by implementing open-source techniques such as third-party “EmptyDrops” (Lun et al., 2019) for cell calling or filtering, which does not assume thresholds specific for Chromium platform.

## Methods

Conceptually, UniverSC carries out its entire process in seven steps (Fig. 1). Given a set of paired-end sequence files in FASTQ format (R1 and R2), a genome reference (as required by Cell Ranger), and the name of the selected technology, UniverSC reformats the whitelist barcodes and sequence files to fit what is expected by Cell Ranger. Additionally, UniverSC also outputs a file with summary statistics including mapping rate, read and UMI counts for each barcode, and reads/UMI for the filtered cells. Sequence trimming based on adapter contamination or sequencing quality is not included in the pipeline and no trimming is required to pass files to UniverSC. However, trimming is highly recommended particularly on R2 files as this generally improves the mapping quality. This requires careful data handling to ensure that all Read 1 and Read 2 are strictly in pairs while only trimming Read 2. We provide a script for convenience that filters Read 1 and Read 2 by quality of Read 2 and avoids mismatching cell barcodes. In principle, UniverSC can be run on any other droplet-based technologies such as inDrop as well as other well-based technologies such as Smart-Seq3 or Quartz-Seq2 (see the software documentation and Table 1 for more details), provided that they contain UMIs. Settings can also be restored to run 10x Genomics Chromium samples as changes made on Cell Ranger by UniverSC are reversible.

**Table 1.** Technologies currently available and settings used by UniverSC.

| Parameter value | Technology | Barcode length<sup>†</sup> | UMI length | Reference |
| --- | --- | --- | --- | --- |
| 10x-v2 (or 10x) | 10x (version 2)<br>Instrument: Chromium<br>Vendor: 10x Genomics | 16 | 10 | Zheng *et al.*, 2017*<sup>6</sup> |
| 10x-v3 (or 10x) | 10x (version 3)<br>Instrument: Chromium<br>Vendor: 10x Genomics | 16 | 12 | 10x Genomics |
| celseq | CEL-Seq | 8 | 4 | Hashimshony *et al.*, 2012;<br>Hashimshony *et al.*, 2016 |
| celseq2*<sup>1</sup> | CEL-Seq2 | 6 | 6 | Hashimshony *et al.*, 2016;<br>Yan, 2017 |
| icell8 | ICELL8<br>Instrument: ICELL8<br>Vendor: Takara Bio | 11 | 14 | Takara Bio Inc.;<br>Goldstein *et al.*, 2017*<sup>6</sup> |
| indrops-v1*<sup>2,3</sup> | inDrop (version 1) | 19 | 6 | Klein *et al.*, 2015;<br>Veres & Lee, 2018 |
| indrops-v2*<sup>2,3</sup> | inDrop (version 2)<br>Vendor: 1CellBio | 19 | 6 | Zilionis *et al.*, 2017;<br>Veres & Lee, 2018 |
| indrops-v3*<sup>3,4</sup> | inDrop (version 3) | 11 | 6 | Zilionis *et al.*, 2017 |
| Nadia<br>(or dropseq) | Nadia or DropSeq<br>Instrument: Nadia<br>Vendor: Dolomite Bio | 12 | 8 | Dolomite Bio;<br>Macosko *et al.*, 2015*<sup>6</sup> |
| marsseq-v1 | MARS-Seq | 6 | 10 | Jaitin *et al.*, 2014 |
| marsseq-v2 | MARS-Seq 2.0 | 7 | 8 | Keren-Shaul *et al.*, 2019 |
| quartz-seq2-1536 | Quartz-Seq2 (1536 wells) | 15 | 8 | Sasagawa *et al.*, 2013 |
| quartz-seq2-384 | Quartz-Seq2 (384 wells) | 14 | 8 | Sasagawa *et al.*, 2013 |
| sciseq*<sup>1,4</sup> | SCI-seq | 10 | 8 | Cao *et al.*, 2017;<br>Vitak *et al.*, 2017 |
| scrbseq | SCRB-Seq,<br>mcSCRB-Seq | 6 | 10 | Soumillon *et al.*, 2014;<br>Bagnoli *et al.*, 2018 |
| seqwell | plexWell<br>vendor: seqWell | 12 | 8 | Gierahn *et al.*, 2017 |
| smartseq | SMART-Seq (version 3)<br>Vendor: Takara Bio (v2) | 11 | 8 | Hagemann-Jensen *et al.*,<br>2019;<br>Ziegenhain, 2020 |
| splitseq*<sup>1,2,5</sup> | SPLiT-Seq | 18 | 10 | Rosenberg *et al.*, 2018;<br>Parekh *et al.*, 2018;<br>Ziegenhain, 2020; |
| surecell*<sup>5</sup> | SureCell<br>Instrument: ddSEQ<br>Vendor: BioRad | 18 | 8 | Bio-Rad;<br>Teichmann Group, 2016;<br>Romagnoli *et al.*, 2018 |

† Barcode length is max or total (linkers are removed automatically where needed) excluding barcodes in the index files which requires demultiplexing.
\*<sup>1</sup> These technologies have their UMIs before their barcodes. The positions of UMIs and barcodes are automatically inverted when these technologies are selected as options.
\*<sup>2</sup> These technologies have their barcodes and UMIs in R2 rather than R1. The functional role of R1 and R2 are automatically inverted when these technologies are selected as options.
\*<sup>3</sup> These technologies have their barcode in two segments with Barcode-1 (8-11 bp) and Barcode-2 (8 bp). The first 19 bp of the adjusted R1 file is originally recognized as barcode, but only the first 16 bp are used upon assigning reads to cells.
\*<sup>4</sup> These technologies have dual indexes (I1 and I2 from the i7 and i5 indexes from Illumina), which contain additional information on cell barcode rather than sample. These require demultiplexing with bcl2fastq before running UniverSC.
\*<sup>5</sup> These technologies have their barcode is in three segments with Barcode-1 (6 bp), Barcode-2 (6 bp), and Barcode-3 (6 bp). The first 18 bp of the adjusted R1 file is originally recognized as barcode, but only the first 16 bp are used upon assigning reads to cells.
\*<sup>6</sup> Test data used in our study was generated from data this paper originally published.

**Figure 1.**
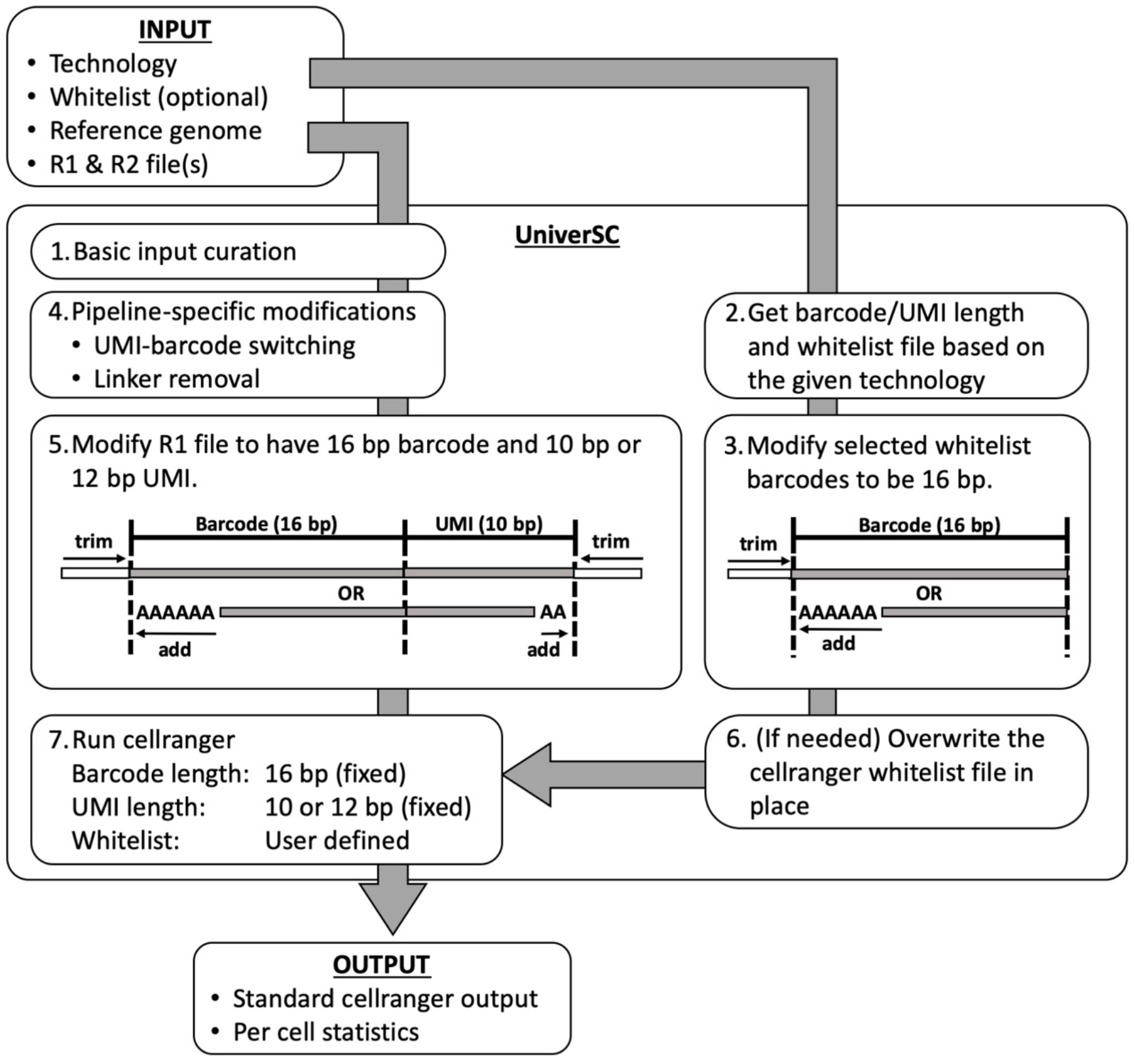
Overview of UniverSC. Given a pair of FASTQ files (R1 and R2), a genome reference (as required by Cell Ranger), and the name of the technology UniverSC first runs a basic input curation (step-1). Then (1) barcode length, (2) UMI length, and (3) the barcode whitelist suited for the technology (if unspecified by the user) is determined (step-2), and the whitelist barcodes are modified to 16 bp (step-3). The curated input files are then adjusted for pipeline-specific modification (step-4) and subsequently reformatted to match the expected barcode and UMI lengths (step-5). If the selected whitelist is different from the whitelist in place for Cell Ranger at the moment, the whitelist is replaced (step-6). Finally, the modified sample data is processed by Cell Ranger against the modified whitelist (step-7) to generate a standard output along with a summary file with per cell statistics.

The set of input parameters for UniverSC is similar to that required by Cell Ranger, with a few additions. The UniverSC workflow requires paired-end FASTQ input files and reference data as prepared by Cell Ranger. By default, UniverSC assumes Read 1 of the FASTQ to contain the cell barcode and UMI and Read 2 to contain the transcript sequences which will be mapped to the reference, as is common in 3’ scRNA-seq protocols. Given a known barcode and UMI length, UniverSC will check the file name and barcodes, altering the configurations to match that of Chromium as needed. The “chemistry” appropriate for each single-cell technology for 3’ scRNA-seq is determined automatically (technologies for 5’ scRNA-seq other than that of Chromium are not supported at time of writing). Data from multiple lanes is supported and so is a custom set of barcodes specific to a given technology other than 10x Genomics.

Test data for 10x Genomics (Zheng et al., 2017) has been downloaded from the 10x Genomics website (10x Genomics, 2020). Test data for DropSeq (Macosko et al. 2015) has been downloaded from GEO (Accession GSE63473) and prepared to match the same reference data. The Nadia technology uses the same barcode design as DropSeq (beads supplied by ChemGenes). Test data for ICELL8 (Goldstein et al., 2017) were obtained from EGA (Accession EGAD00001003443) and filtered for reads matching the same loci.

Each test data was processed in parallel to generate two GBMs consisting of UMI counts. The pair of GBMs were adjusted to have matching sets of barcodes and genes: only barcodes found in both GBMs were kept, and genes only found in one GBM was added to the other with 0 UMIs assigned. The adjusted pair of GBMs were then used to carry out clustering analysis with an R package Seurat. The adjusted pair of GBMs were then used to carry out clustering analysis with an R package Seurat. Finally, the Pearson correlation between the GBMs and the adjusted ARI between the two clustering outcomes were calculated.

## Results and Discussion

We provide documentation for UniverSC accessible as a manual and help system in the terminal and a user-interface which checks file inputs and gives error messages to identify potential problems. UniverSC can be run on any Unix-based system in the shell and the source code is publicly available along with installation instructions at GitHub (https://github.com/minodalab/universc), and a docker image is also available at DockerHub with all dependencies installed from source https://hub.docker.com/repository/docker/tomkellygenetics/universc). We recommend installing UniverSC in a local directory (to a home directory) or somewhere with write access, it can be run on any system with Cell Ranger installed (i.e. added to the PATH environment variable). We also recommend running UniverSC on a server with sufficient memory to run the STAR alignment algorithm. Submission to a cluster in parallel with a job scheduler is supported, but note that UniverSC can only run on one technology at a time due to the different barcode whitelist requirements. A continuous integration service will be used to test updates and maintain the software. See the manual for further details. Note that UniverSC was developed by a third-party unrelated to 10x genomics, and an open-source version of Cell Ranger 3.0.2 is used with Cloupe (a portion of Cell Ranger) inactivated to comply with 10x Genomics End User Software License Agreement.

At initial release, UniverSC has pre-set parameters for 19 technologies (Table 1). Further technologies can be used with “custom” input parameters for any barcode and UMI lengths or by requesting a feature to be added to the GitHub repository. Testing datasets for the following settings are provided: 10x Genomics version 2 and 3 (default), DropSeq, and ICELL8. UniverSC is freely available at GitHub (https://github.com/minodalab/universc), and at DockerHub (https://hub.docker.com/repository/docker/tomkellygenetics/universc). See methods for details on how to install and run UniverSC.

We demonstrate our method using test data from human cell lines that we provide with the package for 10x Genomics version 3 (Zheng et al., 2017), DropSeq (Macosko et al., 2015), and ICELL8 (Goldstein et al., 2017). DropSeq is an example of a droplet-based single-cell technology that does not have known barcodes so a whitelist needs to be generated for compatibility. ICELL8 is a well-based technology that has a known barcode whitelist and allows selecting subsets of wells by known barcodes. These represent two different classes of technologies with different configurations for processing cell barcodes. To assess the degree of similarity between UniverSC and other pipelines, we processed the three test datasets through UniverSC and platform-specific pipeline: Cell Ranger 3.0.2 for 10x Genomics data, dropSeqPipe 0.6 (Roeilli *et al*., 2020) for Nadia data, and CogentAP 1.0 (Takara Bio) for ICELL8 data (Fig. 2). For all three test datasets, the gene-barcode matrix (GBM) generated through UniverSC with the GRCh38 (hg38) was highly correlated with GBM generated by the coupled pipeline (Fig. 2). As expected, it was identical (r=1) in the case of Cell Ranger 3.0.2, and was 0.96 or higher in the two other sets of GBMs. Likewise, clustering results were also highly similar: Adjusted Rand Index (ARI) was 1 in the case of 10x Genomics data, 0.74 and 0.85 for Nadia data and ICELL8 data, respectively. The UniverSC outputs for each of these technologies are provided in the supplementary materials. These results should not be interpreted as biologically meaningful and we recommend running UniverSC on full datasets from experiments or public databases to gain biological insights.

**Figure 2.**
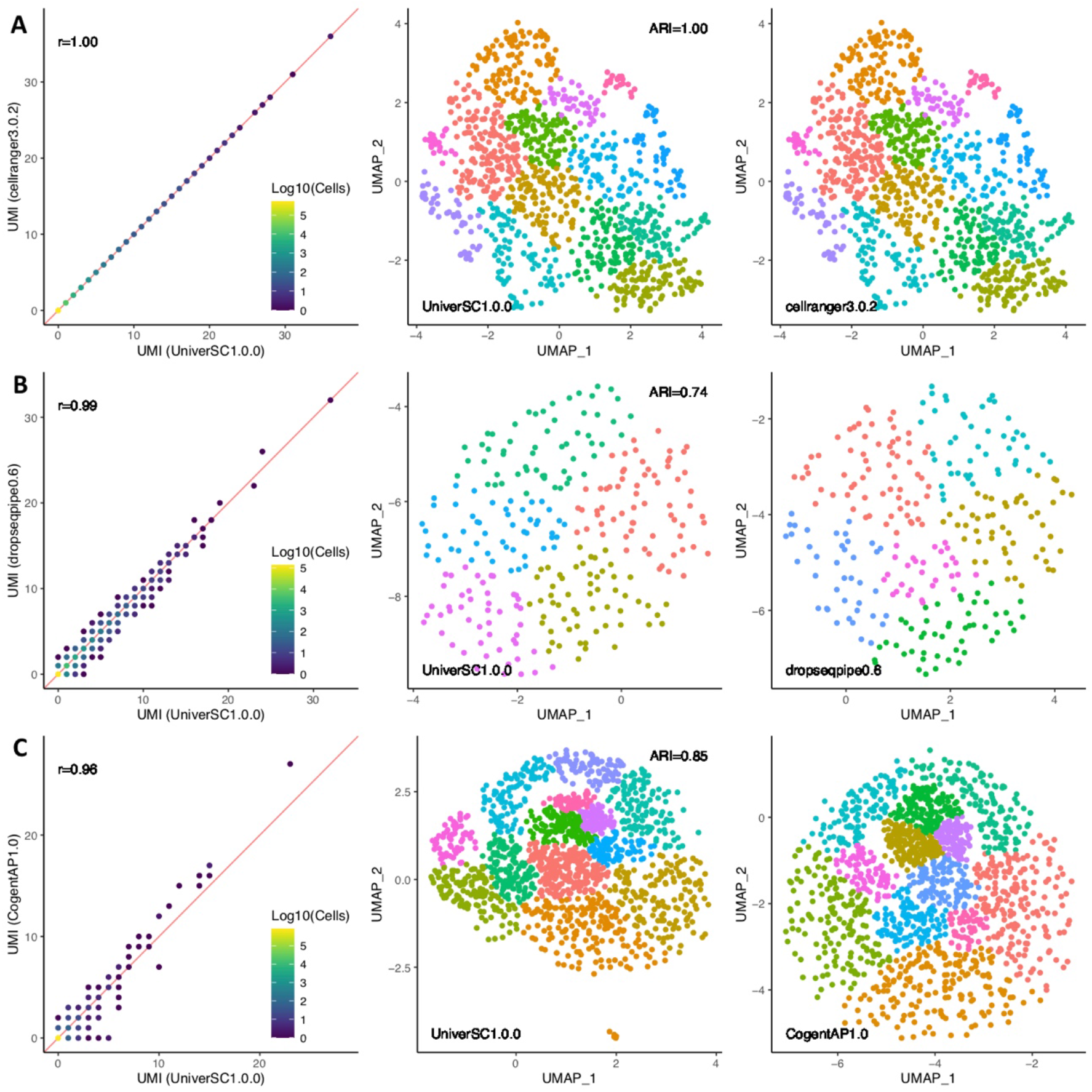
Similarity assessment of UniverSC against other pipelines. Comparisons between the GBM generated by UniverSC 1.0.0 against Cell Ranger 3.0.2 (A), dropSeqPipe 0.6 (B), and CogentAP 1.0 (C). Direct comparison of GBMs is on the left column followed by the clusters resulting from UniverSC 1.0.0 (center column) and its counterpart (right column). The processed data compared here is provided in the supplementary data.

The need for integrating scRNA-seq data generated across different groups and platforms is widely expected to grow as single-cell technologies become increasingly valuable and available in a wide range of studies. Thus, support for processing a wide range of scRNA-seq data with various barcode/UMI configurations under consistent framework is essential. Although there are pipelines that can be configured for a variety of technologies (dropSeqPipe: Roelli et al., 2020; dropEst: Petukhov et al., 2018; Kallisto/BUStools: Melsted at al., 2019), Cell Ranger performs well in a server or cluster environment, and generates a rich and informative output summary. UniverSC enables cross-platform single-cell data integration with a command-line interface, eliminating the need to set up separate pipelines for each platform. We believe our tool will also facilitate the generation of reproducible results. We provide this tool for free and open-source to democratize single-cell analysis for a wide range of scientific applications.

## Acknowledgements

We acknowledge contributions from Tommy Terooatea of RIKEN IMS for testing UniverSC, Jonathon Moody and Chung-Chau Hon of RIKEN IMS for their insightful discussion. We also acknowledge Shuwen Chen, Tsuyoshi Okumo, Max Sanchez, and Karthik Swaminathan (Takara Bio) for support analysing data from ICELL8 platform with their CogentAP pipeline. We thank developers at 10x Genomics of Cell Ranger and dependencies for making their code publicly available. We also thank Marcus Kinsella (CZI) for releasing a docker image of an open-source version of Cell Ranger 2.0.2.

## Glossary

Here follows a summary of definitions used in here and in the package documentation to avoid ambiguity.

### Bioinformatics Procedure

The steps taken to process the data, typically formatting data using existing algorithms and scripting languages.

### Cell Barcode

A short nucleotide sequence incorporated as a part of each fragment in a library that is used to determine the corresponding cell.

### Chemistry

The term used by 10x Genomics to refer to different “versions” of their experimental kits. Here we use it to describe different parameters for Cell Ranger to account for these differences in the chemistry used to prepare the samples.

### Index

The index adapter sequence used in multiplexed sequencing, typically for identifying each sample, using the i7 or i5 indexes for Illumina platforms. Some technologies use these for additional cell barcodes.

### Library

In the context of genomics this refers to the genomic or complementary DNA prepared for sequencing on an NGS sequencing platform.

### Platform

The instrument used to perform single-cell encapsulation (e.g., Chromium, Nadia, ICELL8) or NGS (e.g., HiSeq2500, NovaSeq6000, MGISEQ-2000).

### Single-Cell Encapsulation

An experiment to capture individual cells in wells or droplets.

### Single-Cell RNA-seq

Combining single-cell encapsulation with next-generation sequencing of complementary RNA to gain an expression profile of individual cells.

### Whitelist

The appropriate list of expected barcodes used as reference.

### Unique Molecular Identifier (UMI)

A short nucleotide sequence incorporated as a part of each fragment in a library that is used to determine the corresponding molecule.

